# Replay as a basis for backpropagation through time in the brain

**DOI:** 10.1101/2023.02.23.529770

**Authors:** Huzi Cheng, Joshua W. Brown

## Abstract

How episodic memories are formed in the brain is an outstanding puzzle for the neuroscience community. The brain areas that are critical for episodic learning (e.g., the hippocampus) are characterized by recurrent connectivity and generate frequent offline replay events. The function of the replay events is a subject of active debate. Recurrent connectivity, computational simulations show, enables sequence learning when combined with a suitable learning algorithm such as *Backpropagation through time* (BPTT). BPTT, however, is not biologically plausible. We describe here, for the first time, a biologically plausible variant of BPTT in a reversible recurrent neural network, R2N2, that critically leverages offline-replay to support episodic learning. The model uses forwards and backwards offline replay to transfer information between two recurrent neural networks, a *cache* and a *consolidator*,that perform rapid one-shot learning and statistical learning, respectively. Unlike replay in standard BPTT, this architecture requires no artificial external memory store. This architecture and approach outperform existing solutions and account for the functional significance to hippocampal replay events. We demonstrate the R2N2 network properties using benchmark tests from computer science and simulate the rodent delayed alternation T-maze task.

## 1. Introduction

Forming memories of our lives’ episodes requires the ability to encode and store extended temporal sequences. Those sequences could be things said, places visited or innumerable other sequences of states. Beyond enabling humans’ pastime of recounting our prior experiences, episodic memories are the basis of predictive models of how the world works that support adaptive decision making [1, 2]. How brains build memories of temporal sequences remains poorly understood. It is known that specific brain circuits (e.g., the hippocampal formation[3, 4, 5, 6, 7, 8, 9]) and functional dynamics (e.g., hippocampal replay events) are particularly important. However, the functional principles by which the hippocampus and replay events therein enable sequence / episode encoding remain a puzzle. Machine learning approaches can solve this problem but are biologically implausible. Here, we explore how the principles that underlie machine learning approaches, when modified to be biologically plausible, may elucidate our understanding of how the brain builds episodic memories.

Artificial neural networks, when trained with engineered machine learning approaches, are well capable of encoding protracted temporal sequences. Temporal sequence learning is a task solved particularly well by recurrent neural networks (RNNs). RNNs contain one or more layers of neurons with reciprocal connections among the neurons in that layer (i.e., recurrent connections). This means that the activity of a neuron is a function of both activity in other layers and the activity of its own layer a moment prior. When the recurrent connections are tuned appropriately, the network becomes capable of recognizing sequences, predicting upcoming transitions, and intrinsically recalling sequences. The key challenge, of course, is how to tune the connections. This breaks down to two specific questions. First, ”What learning algorithm allows for reliable encoding and storage of extended sequences from as little as a single experience with that sequence?” and second, ”How can this learning be done in a biologically plausible way?”

A potent learning algorithm that enables artificial RNNs to effectively encode extended temporal sequences is Backpropagation Through Time (BPTT; [10]). Briefly, BPTT works as follows: A sequence of patterns is applied to an input layer of an RNN. A recurrent layer integrates this input along with it own state, generating a temporally evolving pattern of activity. A full record of the spatiotemporal activity of the input and recurrent layers is stored. Given the activity of the recurrent layer, the network can then predict subsequent outputs or any signals that is trained with. Differences between the predicted and actual next states are errors. Errors are the product of the current connection strengths and the past activity of the input and recurrent network layers. Following a presentation to a sequence, during an offline learning phase, BPTT combines information about the connection strengths and activity history while propagating the error back along the computation graph to attribute the blame for the errors to individual network connections (Figure 1A). Finally, individual connections are weakened or strengthened proportionally to their share of the blame for the error. With repeated presentations and epochs of offline learning, the network becomes able to accurately predict or generate the sequence.

**Figure 1:**
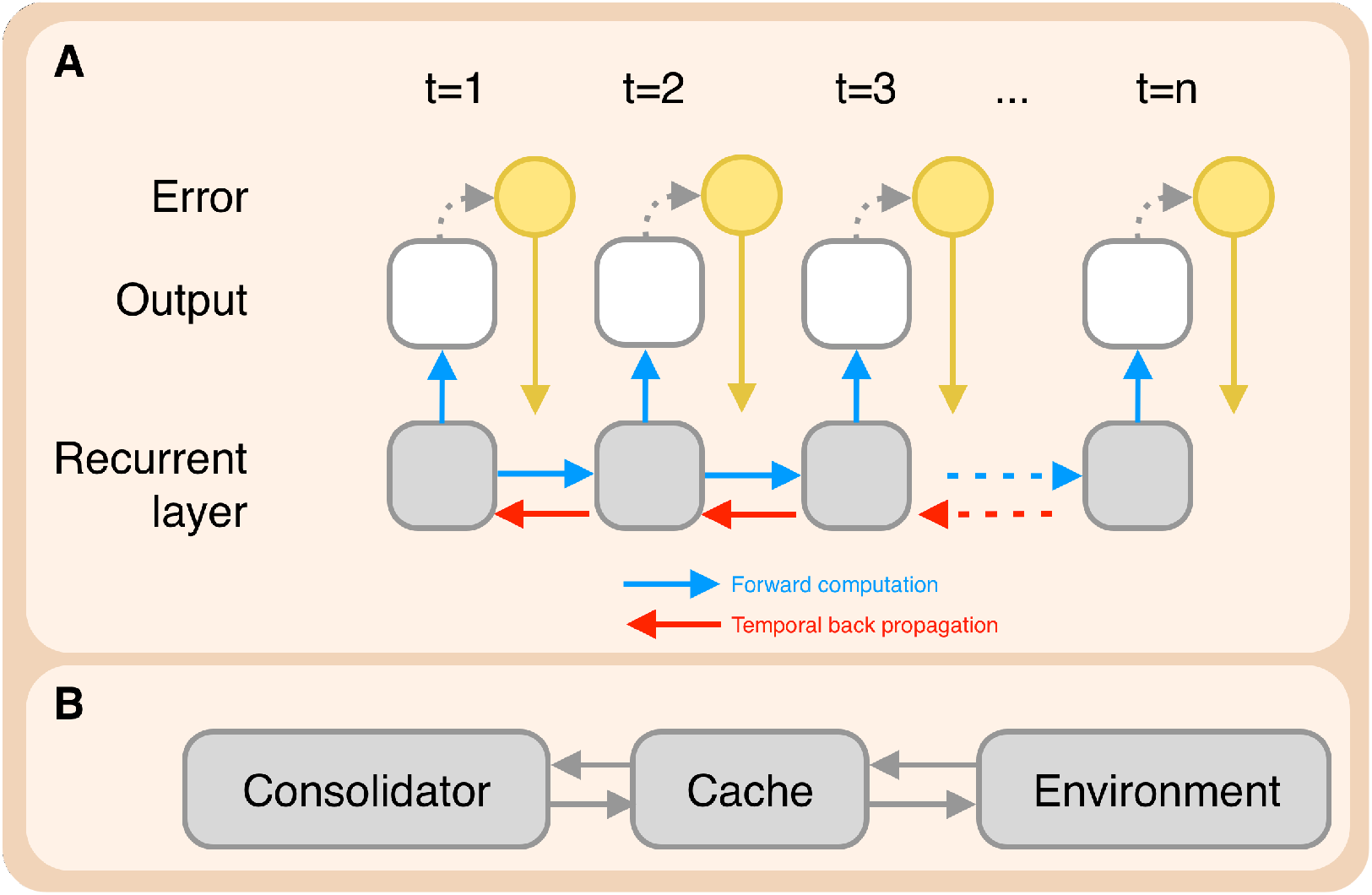
**A** Temporally unfolded computational process of an RNN in BPTT: The recurrent layer at each time step generates an output through the blue projection; error signals (yellow circles) are computed according to the output layer and propagated to the recurrent layer via the yellow projection. To compute the gradient at different time steps, error signals also need to be propagated temporally, i.e. backward in time. **B** consolidator-cache model. The consolidator-cache system first generates output and gets feedback from the environment. Then the cache network stores sequences and plays them back in reverse order to the consolidator network, which in turn optimizes itself based on the replayed content.

Though BPTT is an effective RNN learning algorithm, its value for explaining how brains form episodic memories is questionable because it is not biologically plausible. The implausibility results from violations of the *locality constraint*. The locality constraint captures the fact that biological connections, synapses, can only be changed given locally available information. BPTT violates this constraint in two key ways.

First, BPTT stores and uses an external record of the network’s past activity. Though this activity information was available as it propagated across the network during the presentation of the input sequence, it is no longer locally available during the offline learning phase. Moreover, the prior activation states can not be recomputed with locally available information. This is because the state at time *t* depended upon the state at time *t-1*, information not available at time *t+1*. Ordinary BPTT solves this by literally saving a record of the activation history (e.g., in RAM or GPU memory). Resolving this form of biological implausibility requires addressing how information about the activation history can be obtained in reverse chronological order with only locally available information.

Second, BPTT violates the locality constraint in how the information about current connection strengths is used in the offline learning phase. To attribute blame for error to individual connections, BPTT propagates error backward over the computation graph. This can be accomplished in two conceptually distinct ways but both are biologically implausible. The first is that a separate error network is implemented wherein the connections (e.g., between neurons A and B) are defined as the transpose of the connection in the main network. In other words, the connection B→A in the error network is identical to the strength of A→B in the main network. This makes it so that the error passed from B to A in the error network is proportional to the amount of activity passed from A to B in the main network. In this way, the error network accurately attributes errors to each connection. The main network is then updated accordingly. The key implausibility, called the ”weight transport problem”, is how the connections of the error network mirror those of the main network and how the main network updates are informed by the processing in the error network. Addressing the weight transport problem but suffering from another implausibility is the second way to backpropagate errors. The second approach propagates error backward in the main network itself–firing connections backward with information about the error. This effectively removes the need to transfer connection information between networks but creates the need to pass information backward across connections. Generally, biological synapses do not run in reverse in terms of propagating neural activity [11]. Resolving these implausibilities requires addressing the question that how the connection strength information can be factored into learning so that only locally available information is used.

Establishing biologically plausible means of tuning neural networks to store temporal sequences is an important and ambitious goal. This goal is important for its potential to offer a functional hypothesis for how neural systems support memory for episodes or protracted sequential events. It is ambitious because the native mechanisms of BPTT were engineered to specifically meet the functional needs. Reaching this goal requires addressing biological implausibilities regarding 1) retrieving the activation history of the input and recurrent layers, and 2) how information about connection strengths is used to attribute error across connections and backward in time.

Solutions have been proposed previously for both implausibilities but each has suffered from notable limitations. The utilization of external storage, for example, has been addressed with various approaches. The specifics of those approaches differ but they share a common feature. The solutions omit the need for offline access to a record of the activity through clever handling of the activity while it is still present during the original presentation of the sequence. That is, they compute the sources of error in an online way [12, 13, 14]. These are remarkable for their ability to leverage on-the-fly computations to support learning but these adaptations come at a substantial cost to final performance. Moreover, the omission of offline replay does not improve biological plausibility. Offline forward and reverse replays are well-established to occur in biological neural networks [15]. Indeed, there is strong evidence that offline replay is essential for learning [16]. For offline replay to exist, however, there must be a way to regenerate the patterns. Defining how this occurs is a puzzle that we address in the present work.

Solutions also exist to address the weight transport problem and reversible connections problem, but they too have limitations. For example, the reversible connection problem comes up in training feed-forward networks. In that setting, it was shown that knowledge of the weights is not needed and that fixed random top-down connections can function to train various networks [17] (see also [18]). This is referred to as *feedback alignment*. Though originally designed for feed-forward networks, feedback alignment can also be used in RNNs. One such variant is called *random feedback local online (RFLO)* [13]. The feedback alignment approach of RFLO effectively addresses the biological implausibility issue. Critically, however, RFLO is functionally limited to propagating error only one step backward in time [19]. A method capable of tracking the temporal gradient over many time steps in RNNs remains lacking but is a gap we address in the present work.

We present here a biologically plausible recurrent network model of episodic learning based on BPTT without suffering from the biological implausibilities of BPTT. This model, referred to here as R2N2 (short for *Reversible Recurrent Neural Network*), fully satisfies the locality constraint. R2N2 uses no external record of the network’s past activity. Instead, it leverages two previously described separate solutions for enabling reversible reactivation of a network, one for the input layer and one for the recurrent layer. Further, R2N2 neither transports weights nor does it assume reversible synapses. Instead, it leverages an error network that is controlled by and controls the main network in a way that allows error backpropagation to train the network without weight transport. The individual components of R2N2 are each based on established approaches. The full R2N2 model, and the fact that it collectively represents a high-functioning biologically plausible replacement for BPTT, is novel and innovative. The specifics of each separated solution and the operation of the full R2N2 model are described in the Model section. In the Results section, we demonstrate the sufficiency of each solution separately. We then combine the components to form R2N2 and benchmark the performance of this fully biologically plausible implementation of BPTT, showing it surpasses current state-of-the-art biologically plausible implementations. Finally, to facilitate comparison to sequence learning in brains, we show that R2N2 can learn the classic delayed alternation T-maze task. While our model is designed to replace biologically implausible components with ones that are plausible in principle, it is not designed to simulate specific anatomy or physiology. Nonetheless, as illustrated and discussed, the full model recapitulates several key functional properties of the hippocampal formation including place cells and offline replay.

## 2. Model

The full model consists of two interconnected RNNs referred to here as the consolidator and the cache (Figure 1B). We refer to the full model as the *Reversible Recurrent Neural Network* or R2N2 for short. The consolidator and cache are both RNNs but with different architectures and learning rules as each is designed for distinct functions. These are briefly summarized here, and specifics are unpacked in detail in the sections below. The consolidator is the primary RNN, designed to have a large storage capacity and robust generalization ability. A trained consolidator is functionally akin to an RNN trained with BPTT. The cache is an auxiliary network, designed to support the training of the consolidator. The cache is optimized for rapid encoding and high-precision bi-directional retrieval of input sequences. This enables the cache to perform one-shot learning of to-be-learned sequences. During offline processing, the cache replays the sequences in reverse order to the consolidator to train the durable trace of the memory.

### 2.1. The Consolidator Network

Taking inspiration from [20], we developed the consolidator network, an RNN that is composed of at least two interconnected populations of neurons A and B (see Figure 2A). The neuron group A receives input signals from neuron group B and itself and vice versa. More specifically, we divide the incoming projections for each group into two parts, and the corresponding dynamics equations can be written as follow:

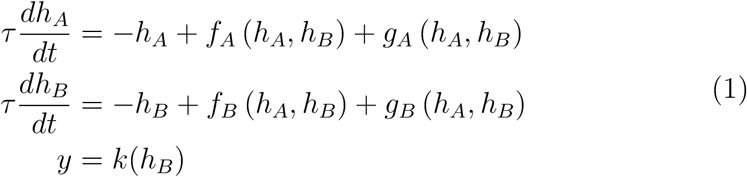

We define the *f*_∗_ (∗ can be *A* or *B*) projections as forward connections with the *g*_∗_ being the backward projections and *k* being the output projection. In this part, we will be focused on the recurrent units in A and B and leave the discussion of output *y* to later parts, in which the network firing rate dynamics generated by *f*_∗_ is 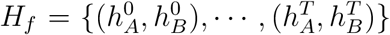, which is called the forward sequence and represents the normal running or forward replay phase of this RNN. To generate a reversed sequence of *H*_*f*_, mathematically, one needs to simply flip the sign of derivatives in the dynamics equation from *t* = 0 to *t* = *T*. If we discretize the dynamics equations into a different form, it will degenerate to the case of a reversible deep neural network block [20], which has been shown to be memory-constant in various tasks as the storage requirement of neural activity equals the number of units in the network.

**Figure 2:**
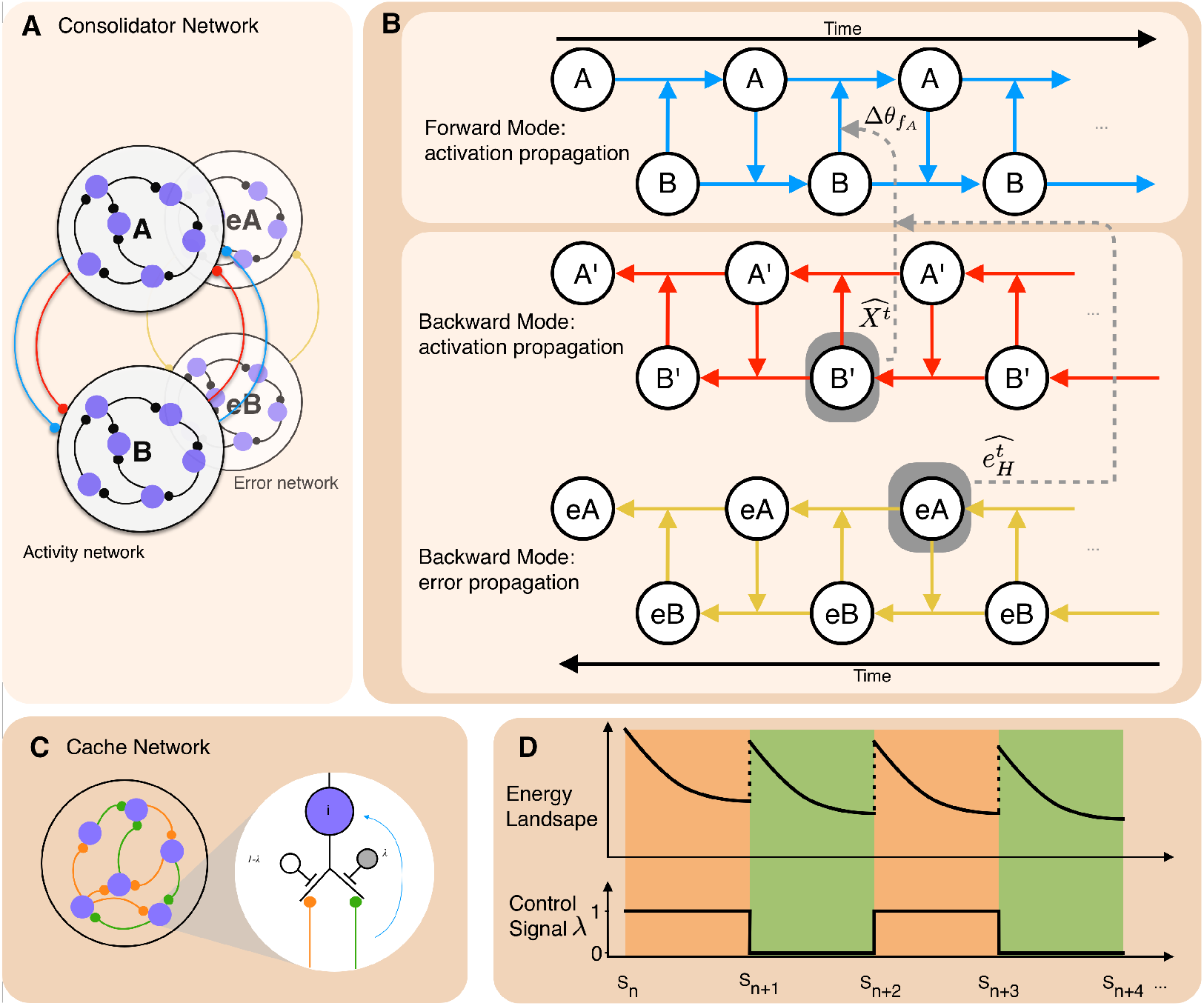
Schematic structure of consolidator and cache. **A** Projections of neuronal group A and B in the consolidator network. The consolidator network is composed of an activity network and an error network. The activity network shown in red represents *f*_∗_ while the network in blue represents *g*_∗_. Notice that the direction of sequence play (forward or reverse) is not fixed by the arrow but determined by the competition between *f*_∗_ and *m, g*_∗_. **B** The temporal unfolding of forward and backward computation in the consolidator. When projection *f*_∗_ is stronger, the whole network operates in the forward mode. In this case, the network updates its activations in both group A and B, generates outputs, and computes error signals accordingly. When the *g*_∗_ connection is stronger, the whole network turns into backward mode. The neuronal group A and B generate reversed activation sequence and thus can be utilized to propagate activation backwards to reconstruct the previous time step activities, bypassing the temporal credit assignment issue caused by non-locality and obviating the need to store previous activities. The error network recursively multiplies the error vector by the feedback alignment random matrix to compute the error vector at the previous timestep. **C** Projections in the cache. In the cache, each neuron receives projections from both *W*_*E*_ (orange) and *W*_*O*_ (green) (**C** left). A detailed description of connection pattern in a single neuron *i* can be found in C right.The final incoming weight of neuron *i* is determined by λ ∈ (0, 1), which can be viewed as a result of competing oscillating interneurons tuning inputs from *W*_*E*_ and *W*_*O*_ synaptic inputs. **D** A schematic description of state transitions in cache. By periodically switching between *W*_*O*_ and *W*_*E*_ via the control signal λ (lower panel), the network operates in two set of weights. Each of them builds state attractors between successive states *S*_*n*_ and *S*_*n*+1_(energy landscape slopes between two edges of the same phase). Different successive state pairs are connected in a chaining way and thus form a long states sequence (upper panel).

The intuition is depicted in Figure 2B. The blue circuit represents a transformation ⊕ (⊕ represents the operation that combines *A* and *B* together) from *A*^*t*^, *B*^*t*^ to *A*^*t*+1^, *B*^*t*+1^ in the next time step: *A*^*t*^ ⊕ *B*^*t*^ → *A*^*t*+1^ and *B*^*t*^ ⊕ *A*^*t*+1^ → *B*^*t*+1^. Then, as long as there exist another inverse operation ⊖ that satisfies *X* ⊕ *Y* = *Z* ⇔ *Z* ⊖ *Y* = *X*, we can construct a circuit that turns *A*^*t*+1^, *B*^*t*+1^ into *A*^*t*^, *B*^*t*^: *B*^*t*+1^ ⊖ *A*^*t*+1^ → *B*^*t*^ and *A*^*t*+1^ ⊖ *B*^*t*^ → *A*^*t*^.

In effect, these two operators (⊕ and ⊖) reverse the update process without requiring any other constraints: the two variables *A* and *B* could be either scalar (signal neuron case) or high dimensional vector (neuron group case). One of the simplest pairs of operators that meets the above conditions consists of addition and subtraction, which is the operator set used by [20]. To map these two operators to the brain seems implausible as it requires flipping the excitability of a synapse in a short time range. However, we can approximately reach the same effect by introducing competition from another group of backward projections *g*_∗_ (∗ can be *A* or *B*). To function, the *g*_∗_ projections must generate currents that have the same amplitude but with different signs of *f*_∗_ to cancel *f*_∗_ − *h*_∗_: *f*_∗_ − *h*_∗_ +*g*_∗_ = 0. This can be easily implemented with local dendritic propagation and local training (see Eq. 2) on *g*_∗_’s parameter *θ*_*g*∗_. This detailed balance between *f*_∗_ and *g*_∗_ projections thus makes it possible to run the whole system in a backward manner: If the excitability of a trained *g*_∗_ projection is scaled by a factor of 2, the resulting effective projection will be approximately *h*_∗_ − *f*_∗_.

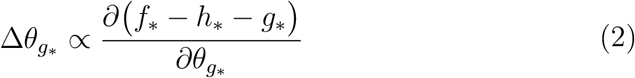

There are many possible choices for *f*_∗_ and *g*_∗_ projections. For example, each can be a simple two-layer neural network if we consider the branching structure of dendrites, which has been shown experimentally to be equivalent to multi-layer nonlinear neural networks [21]. In this case, a complex *f*_∗_ − *h*_∗_ architecture can be approximated by *g*_∗_ as long as the complexity of *f*_∗_ is not higher than *g*_∗_. For demonstration, in the following experiments, we choose the simplest form of *f*_∗_ to be a single-layer nonlinear network and *g*_∗_ to be a combination of one-layer linear and nonlinear networks. Mathematically, this reduces the learning process of backward projection *g*_∗_ to be a regression process, which is known to be trivial to solve with local learning rules but is still enough to support complex sequential computation as non-linearity is involved in each time step. This architecture (Figure 2A) is scalable and thus it is possible to support more complicated computations. If one adds another Group along with *A* and *B*, the coupling can be extended, and the backward running can still be preserved.

Figure 2A depicts a network that can produce sequences of activity in reverse, and the same network structure can also be used to propagate error signals in reverse. Figure 2B shows two network graphs, one which runs activity sequences in reverse, and one which runs error sequences in reverse.

In BPTT (Eq. 3), the error signal 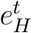 used to update hidden layer connections 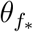 are recursively computed in a mirroring circuit (yellow projections in Figures 2A and 2B) of the forward circuit from ”future” to ”past”, i.e., 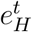 relies on both the future hidden layer error 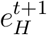 and the transient output error *L*^*t*^.

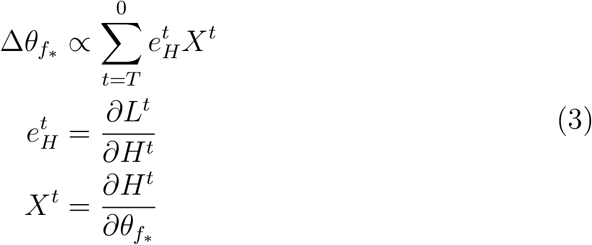

Two types of implausibility exist in this process: 1) the weight transport problem, i.e., how to compute the transient component of 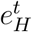 with *L*^*t*^ [22] and 2) the external storage of activations, i.e., how to compute *X*_*t*_ in Eq. 3. On the one hand, for the first issue, [17] proposed an alternative local error circuit, Feedback Alignment (FA). With fixed random projections, it has been shown to be effective on various deep network architectures and tasks [23, 24]. On the other hand, our synaptic competition balance mechanism described above addresses the second issue, as it reconstructs the previous network states *X*_*t*_ in a backward manner. This eliminates the need to store the neural activities at multiple time steps in the past.

Together these two mechanisms propagate the error temporally back with a simple linear tuning of projection excitability and without any non-local information (the first line in Eq.3). The FA algorithm approximates hidden layer errors 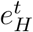 in Eq.3 with 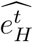 at each time step using *L*^*t*^ and a fixed random feedback matrix. For the specific derivation of 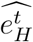 and its recursive update equation, see the supplementary material. The backward projections (red connections in Figure. 2A) in the consolidator network replace *X*_*t*_ with 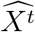 by approximating *H*^*t*^ with 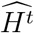. i.e., the reconstructed neural activities generated in a reverse replay. The product of 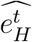 and 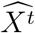 is then used to update forward projections *θ*_*f*∗_.

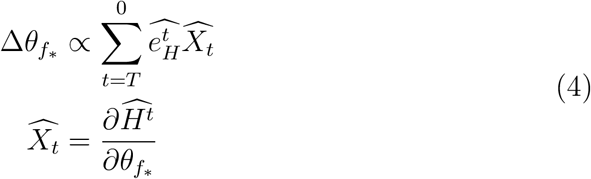

### 2.2. The Cache Network

The learning mechanism described so far (Eq.3) is effective when the consolidator network is solely determined by its previous states, i.e., without external inputs, as the reverse replay equation will not hold if we add a time-varying term in Eq.1. This limits the use cases of the consolidator as most of the sequence learning tasks involve dealing with temporal inputs. To perform reverse replay in a running consolidator that integrates time-varying input sequences *I* (see Eq. 5) through the mapping *b*, external storage of sensory sequence inputs *I* becomes necessary, so that Eq 1 is modified as:

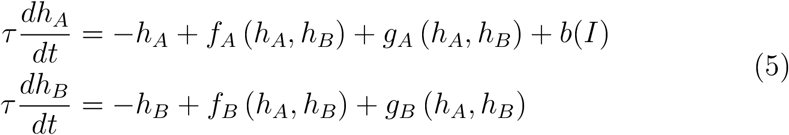

In the replay phase, the consolidator itself cannot generate the dynamics without knowing *b* and, by extension, *I*. Superficially, this brings us back to the original dilemma, i.e., to design another RNN that can run backward. The difference is that it should have the capability to memorize a given sequence after as little as a single exposure, which makes the problem harder. Nevertheless, the fact is that sensory input sequences in most cases are usually in a space that has much fewer dimensions compared with the number of neurons in consolidator, and this suggests a solution.

To memorize sequential sensory inputs and play them in a reversed manner, one can build point attractors representing inputs in the state space and connect them with directed line attractors (Figure 2D). A modified Hopfield RNN (Figure 2C) matches these desired characteristics. A classical Hopfield network builds energy basins that allow noisy inputs to settle into corresponding attractors. A modification to its learning rule, from Δ*W* ∝ *I*·*I*^T^ to Eq.6, then links one attractor to another (*I*^*t*^→*I*^*t*+1^) in the state space with a direction pointing to *I*^*i*^ if this trial is rewarded (*r* = 1). This is the general Hopfield weight update equation for the cache network:

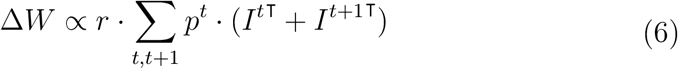

By linking multiple (*I*^*t*^, *I*^*t*+1^) pairs from *t* = 0 to *t* = *T* with the modified learning rule, a reversed pattern sequence {*I*^*T*^, *I*^*T* −1^,, *I*^0^} is built.

The cache is an RNN using the above learning rule combined with time-varying weights [25] to increase the stability of transitions between successive sensory patterns (Eq.7). Once *I*^*t*^ is stably transformed to *I*^*t*−1^ through *W*_*E*_, another group of weights *W*_*O*_ will dominate the transition from *I*^*t*−1^ to *I*^*t*−2^ through the tuning of *λ* (see in Figure 2C the yellow and green projections tuned by two competing interneurons), which can be viewed as an external periodic control signal or a signal indicating the stability of cache activations [26]. In terms of a physical analogy, one can imagine this as a reciprocating pump, in which *W*_*E*_ drives the system from *I*^*t*^ to *I*^*t*−1^, and then *W*_*O*_ drives the system from *I*^*t*−1^ to *I*^*t*−2^, and so on back and forth between *W*_*E*_ and *W*_*O*_, by the following equation that governs the activity of the cache network:

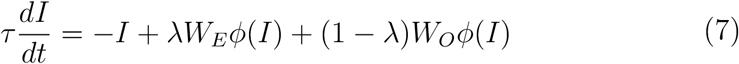

The cache network thus can learn arbitrary sequences of patterns in a local, stable, and one-shot manner as the weights update rule of the Hopfield network is local and can be computed with only a single exposure to the inputs.

### 2.3. R2N2: Sequence learning with consolidator and cache

In training a vanilla RNN with BPTT, one needs to perform the following steps:

- Initialize the RNN and run forward with temporally varying inputs.
- Store the inputs sequence, hidden unit activations, output sequence generated and the target sequence to an external memory device.
- After the whole input sequence has been received, compute the error between output and target for the last time step.
- Extract input, output and target pairs from the memory device in a temporally reversed order, propagate the error in a backward manner and compute weight changes simultaneously.
- Apply the accumulated weight changes after finished the backward running phase.

The first point to note here is that the external storage is where the main biological implausibility lies. It’s unclear how the brain could store the activity in each cell at each timestep somewhere else and replay it precisely. However, with the consolidator and cache, this activity memory can be reconstructed dynamically. Notably, the storage size requirement is substantially reduced, as the consolidator can reproduce its historical activations as a reverse-play sequence with the help of the cache. Consider a case in which consolidator has 128 neurons and the channel size of inputs is 16. The standard BPTT needs to store a sequence of 16+128 = 144-dimensional vectors as all input and hidden state vectors need to be preserved in the temporal unfolding process. However, in our model, the system only requires cache to store a sequence of 16-d vectors representing only the sequence of input vectors, because the consolidator can reconstruct its activity by itself. This means a memory of sensory experience instead of all neural activities is enough to support sequence learning. This approach also matches the empirical findings that the replay of location sequences improves animals’ performance in given spatial navigation tasks [27].

Secondly, we modify the standard learning process in BPTT to fit our model. In BPTT, the input and target channels usually belong to different categories. Taking the classical random dots perceptual decision-making task as an example, the input is usually set to the coherence of randomly moving dots’ directions and the desired output target is the eye motion direction [28]. This makes the backward running phase more complicated as the system needs to store the desired target and input patterns together and only compute the error signal based on the difference between generated outputs and desired targets. Instead, the process can be simplified if there is no categorical difference between desired outputs and inputs. Taking inspiration from predictive coding in sequence learning [29], we view performing cognitive tasks as a process of online sequence prediction: the task-relevant stimuli, action signal, and reward signal are treated equally and are concatenated into an integrated ”sensory inputs” vector. Regardless of their structure, various cognitive tasks then can be reduced to the same type of sequence prediction task. Thus the task reduces to predicting the future state at time *t* + 1 on the basis of task-relevant variables at time *t*.

Based on these assumptions and modifications, we propose that learning a specific task can be divided into two phases, with the first one mapped to fast learning and the second one to slower statistical learning, essentially as a consolidation process. In the first stage, the animal explores the task settings and environment randomly, generating both rewarded and unrewarded ”sensory sequences” involving all task-relevant variables. During this initial phase, the cache memorizes ”sensory sequences” that are rewarded at the end of each trial, which can be learned in a one-shot fashion as it is a Hopfield network in principle (see Eq.6).

In the second phase, the cache starts reverse replay, sending signals to the consolidator and thus training it. This means a target for the cache at time *t* is actually input for both the consolidator and cache at time *t*+1, so that the cache does not need to store a target sequence separately. Then the consolidator in the second phase optimizes its forward projections *f*_∗_ according to the targets provided by cache and its own reconstructed reversed activations. Once its forward projections are changed, the backward projections *g*_∗_ will be adjusted accordingly to cancel *f*_∗_. Notice that the adjustment of *f*_∗_ and *g*_∗_ (Eq.4 and Eq.2) could occur simultaneously as the learning of backward projection is an online process. Consequently, the knowledge about the rewarded sensory experience is transferred from cache to consolidator via fast learning at first and then statistical learning later. Besides, as the consolidator can go back to states it experienced, it can also perform forward replay using projections *f*_∗_, which could be used to explore possible future outcomes when the model is in an intermediate state [30, 31]. To sum up, we view this process as an implementation of Buzsaki’s two-phase model [32] for training long-term memories as the interplay between consolidator and cache in two phases simulates the entorhinal-hippocampal communication.

## 3. Results

### 3.1. Consolidator

The consolidator is the primary long-term memory store of R2N2. The architecture is illustrated in Figure 2A and 2B and described in Section 2.1. To be a biologically plausible high-functioning network, the consolidator must demonstrate two capabilities with only local information: 1) *Reversibility:* The ability to reconstruct its activity backward; and 2) *Error backpropagation:* The ability to tune connection proportionately to the errors produced so that error can be reduced. BPTT uses non-local solutions to achieve both capabilities. In this section, we show functioning solutions to both that use only local information. Competing complementary subnetworks[20] can reconstruct the spatiotemporal consolidator activity patterns in reverse without an external record. Feedback alignment[17] can reduce the reconstruction error over training without weight transport or reversible synapses. It should also outperform the highest functioning biologically plausible algorithms. We benchmark R2N2 performance against BPTT and demonstrate that R2N2 performs better than RFLO and echo state networks.

#### 3.1.1. Reversibility

To achieve reversibility in the consolidator network, we used the competing subnetwork approach described previously[20]. To demonstrate reversibility, the consolidator should be able to reconstruct an activation sequence in reversed temporal order without external signals so that the error can be aligned to the dynamics that led to the error.

A random time-varying pattern of activity was applied to the input layer of the consolidator that induced a complex time-varying pattern of consolidator activity. A representative example of the consolidator activity is shown for five neurons in each of the consolidator subnetworks in Figure 3A. The activity of each consolidator subnetwork is, in part, a function of the input from the other subnetwork by way of connections *f*_∗_ (see Model for implementation details). A separate set of inter-subnetwork connections, *g*_∗_, learn to be equal and opposite in sign to *f*_∗_ through a local learning rule (i.e., without use of non-local information). It is proper training of *g*_∗_ that allows for reversible reconstruction of the consolidator activity. Figure 3B illustrates the reversibility of the consolidator after 4 epochs of training. Shown are 100-time steps of the same five neurons as shown in 3A as the newly trained *g*_∗_ connections control the consolidator activity. Comparing this activity to the forward pattern by flipping the time axis and subtracting it from the forward pattern reveals that the activity was well matched, as shown in Figure 3C.

**Figure 3:**
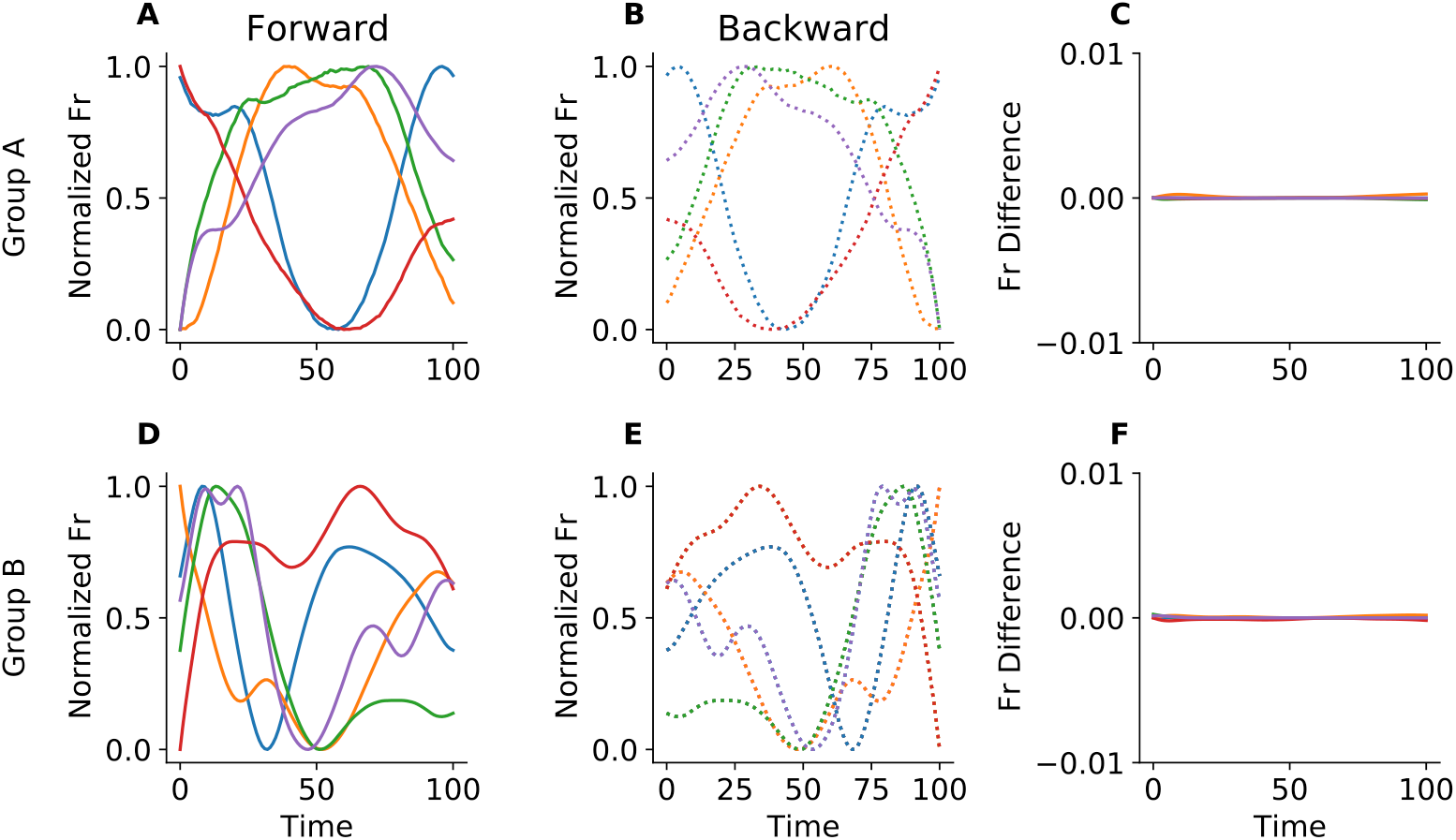
Normalized firing rates of neuronal group A and B in the consolidator (For each group, the first 5 neurons are selected, while overall the consolidator has 128 neurons). **A**,**D:** Solid lines represent firing rates in forward running **B**,**E:** Dashed lines represent firing rates during backward running. **C**,**F:** Difference of firing rates between forward and backward running (flipped). Signs of symmetry are shown clearly in comparisons between the **A** and **B** pair and **D** and **E** pair.

#### 3.1.2. Error Backpropagation with Feedback Alignment

Backpropagation allows a network to take error information that becomes available at the end of a sequence and retroactively tune connection strengths to reduce error. Successful backpropagation requires both a record of the prior activity and a means of relaying the error signal. The record of prior activity in the consolidator is provided by the reversibility property shown above. To relay the error signal, we used the feedback alignment approach described previously[17]. To demonstrate successful backpropagation, the consolidator should be able to adjust its connections to be able to minimize error and thereby reconstruct an input sequence.

A random binary vector data stream was generated to serve as inputs as shown in Figure 4A. Notably, the random inputs included repeated elements at both adjacent and remote time points challenging the network to attribute the error appropriately as a function of time (i.e., not simply learn that state Y always follows X). The consolidator was trained to generate input patterns in the next time step (i.e., predict transitions) using data at the current time step as a cue. After 50,000 training steps, the consolidator was able to predict the random binary vector as shown in Figure 4B. This experiment shows that the consolidator can learn to map input patterns to outputs at a given time despite using shared weights across multiple time steps. This implies that the temporal credit assignment problem is solved effectively (i.e., the correct connections were adjusted for an error resulting from an earlier activity pattern) through the use of feedback alignment and consolidator reversibility.

**Figure 4:**
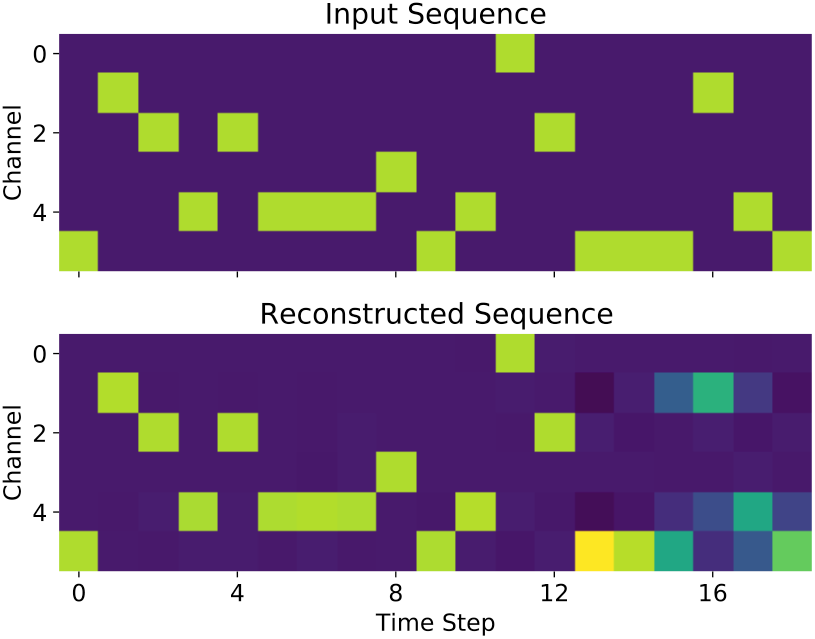
Sequence memorization task for the consolidator. The consolidator is trained to recall elements in the next step using data from the current time step. The top row represents the input data stream and the bottom row represents the sequence generated by the consolidator.

#### 3.1.3. Performance

To benchmark the consolidator performance, we compared the memorization capacity of the consolidator to BPTT and two high-performing, biologically plausible, sequence encoders, Random Feedback Local Learning (RFLO)[13] and Echo State Networks (ESN)[33].

We compared the performance of the four algorithms (consolidator, BPTT, RFLO, and ESN) on a character prediction task *a*^*n*^*b*^*n*^ (Figure 5A). The *a*^*n*^*b*^*n*^ character prediction task[34] is a classic test for assessing RNN capability to encode sequences in the face of strong interference. In short, the input stimuli is a stream of *n a*s and *n b*s (see *Model* section for details). Performing this task requires accurately predicting whether the next character is another repeat or a switch and this requires staying oriented to how many repeats have come already.

**Figure 5:**
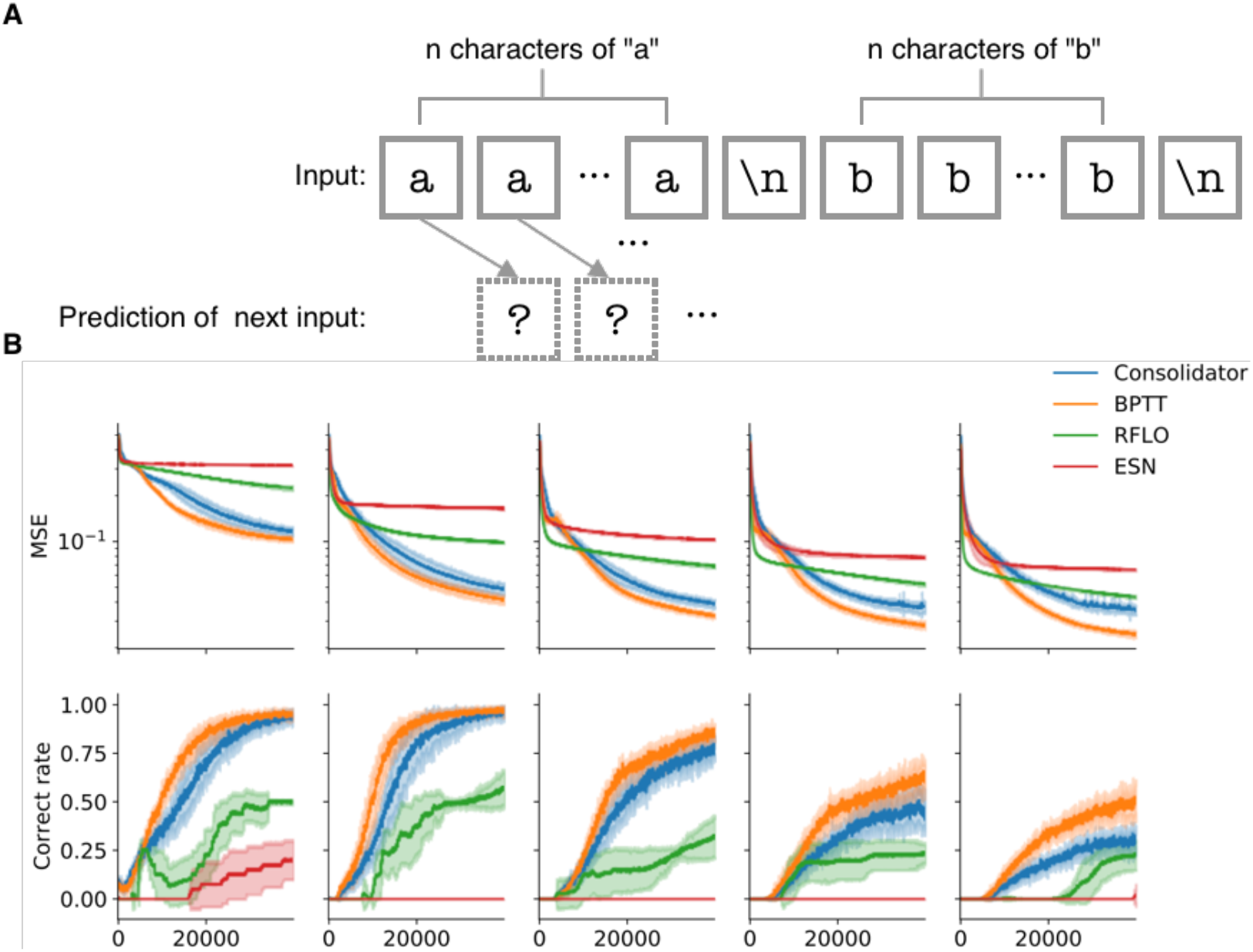
**A** A schematic view of *a*^*n*^*b*^*n*^ task. Each model is trained to predict the next character based on the previous inputs. **B** Performance Comparison on the *a*^*n*^*b*^*n*^ task. The upper rows show averaged loss curves while the lower rows show prediction accuracies. From the leftmost to the rightmost column, the range of *n* in each dataset is linearly increased from a bin including (1, 4) to a bin including (21, 24), thus the corresponding sequence length ranges from (4, 10) to (44, 50). Orange lines represent the consolidator with transpose of forward matrices as backward matrices. Blue lines represent consolidator with fixed random backward matrices, implementing feedback alignment. Green lines represent RFLO, and red lines represent ESN.

The panels of Figure 5B show the mean squared error (MSE) and correct rates of the consolidator (blue), BPTT (orange), RFLO (green), and ESN (red) for sequences of increasing length. The leftmost panels show the performance for input sequences where *n* = 1, 2, 3, 4 and the rightmost panels show the performance when *n* = 21, 22, 23, 24. Given differences in how each algorithm handles the temporal gradient (as unpacked in the *Model* section), we expected that the consolidator performance would be close to BPTT and better than RFLO (Performance: BPTT ≥ consolidator *>* RFLO *>* ESN). The results match our prediction and are shown in Figure 5B. Across sequence lengths, the consolidator performs consistently better than RFLO and ESN, approaching the performance of the biologically implausible BPTT.

Trained RFLO networks can reach a correct rate with an upper bound at around 0.5, which reflects that the error gradient in RFLO is essentially limited to one step backward in time [19], while the consolidator can propagate the error gradient backward multiple steps in time. By comparison, the ESN can hardly generate any proper outputs when the sequence length is longer than 10 (right four columns in Figure 5B). This shows the advantage of multi-time-step temporal error signal propagation over online error minimization, even with the constraint of no external storage of neural activation history.

For shorter sequences (e.g., n *<* 9), the consolidator and BPTT have similar asymptotic performance. As the sequence length increases, the performance of all models decreases (bottom row of Figure 5B), and the divergence between a consolidator implemented with BPTT (orange lines) versus Feed-back Alignment (blue lines) gradually increases. This divergence shows that feedback alignment does have limits relative to pure BPTT, which maintains a perfect record, for temporal propagation across numerous time steps.

### 3.2. Cache

The cache network functions as the primary input to the consolidator. Functionally, it performs rapid memorization of the input sequence for subsequent playback to the consolidator during the offline learning phase. BPTT uses an externally stored record of the input sequence that is aligned with the backpropagated error. To be a biologically plausible high-functioning network, the network must be capable of storing a sequence of states in a way that can be retrieved in reverse order (to synchronize with the reverse replay in the consolidator) after a single training trial.

To achieve this, the cache is itself a classic form of a recurrent neural network, using well-established learning principles (akin to a Hopfield network with multiple weight matrices) that enable retrieval of stored states in forward or reverse order as described in full detail in the Model section. To demonstrate this ability, we tested the ability of an isolated cache network (i.e., with no consolidator network connected) to retrieve a sequence of randomly generated binary vectors.

As shown in Figure 6, the cache was presented with 20 distinct binary vectors over time. Transitions between adjacent vectors were encoded by alternating weight matrices based on a control signal (Figure 6A). With only the single presentation, the cache can step through the same set of states (Figure 6B). This playback can be performed in the forward or backward direction depending upon which pattern the cache is initialized with. The timing of each transition is controlled by the control signal, allowing the network to intrinsically regulate the retrieval. With this ability, the cache can support playback of the input sequence so that it is synchronized with consolidator processing.

**Figure 6:**
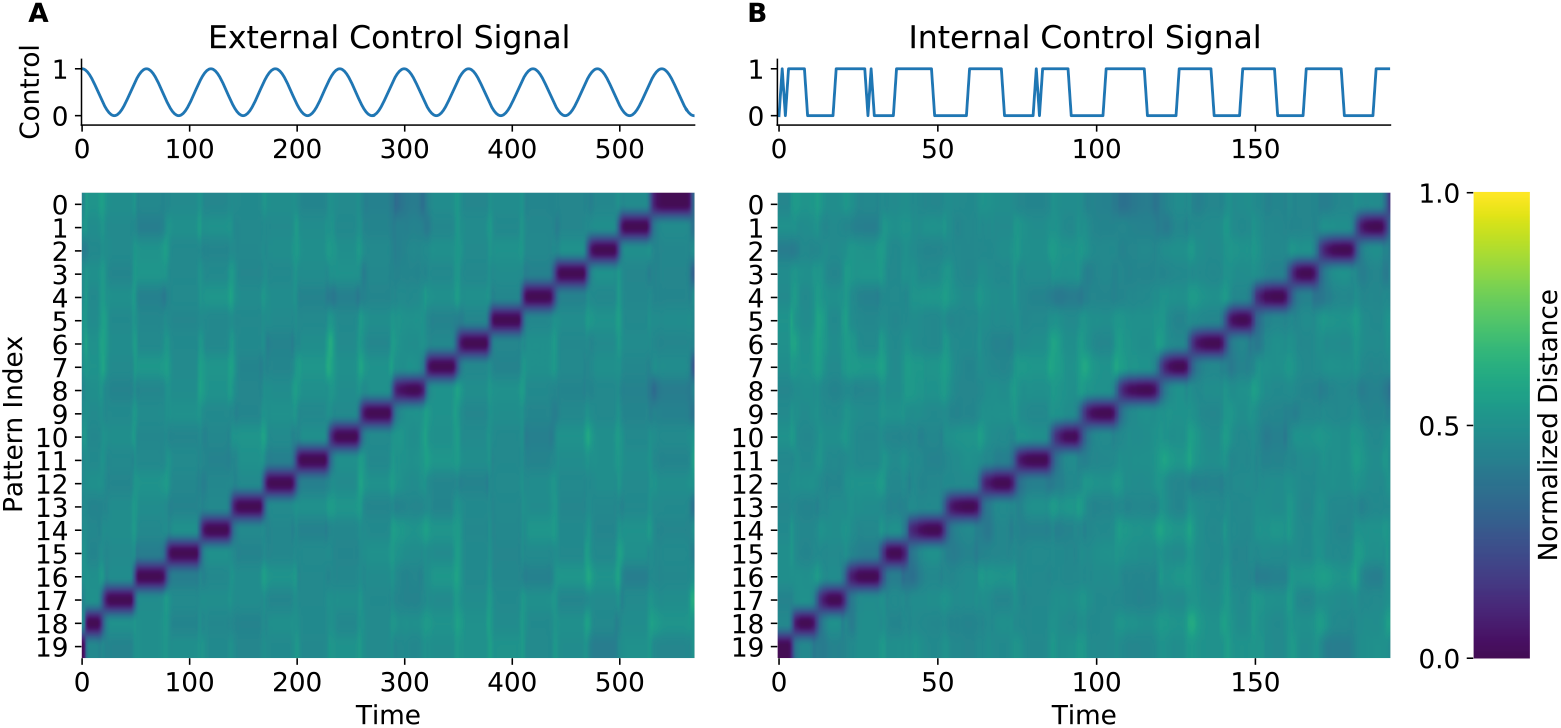
One shot sequence memorization task for the cache. **A** A cache trained to recall the random binary sequence with an external periodic signal. Top: the oscillating external control signal λ, Bottom: The relative hamming distance between cache’s activation and all patterns. A larger pattern index represents a pattern that appears later in the given sequence. **B** Same as **A**, except that it uses an internally generated control signal.

### 3.3. R2N2 Solving Sequence Learning Problems

The results shown thus far demonstrate that each component of R2N2 is capable of performing its function as intended. In this section, we demonstrate that nothing is lost and nothing additional is needed when the components are assembled into the full R2N2 model while adhering to the locality constraint. That is, we show that R2N2 is capable of encoding the memory of temporal sequence into a recurrent neural network using only local information. Given that our motivation was to understand how brains enable episodic memory, we applied R2N2 to a simulation of a T-maze navigation task, in which the animal needs to make decisions to either turn to the left or right end of the horizontal branch to get a reward based on the visual cue at the beginning of the maze (see Figure 7).

**Figure 7:**
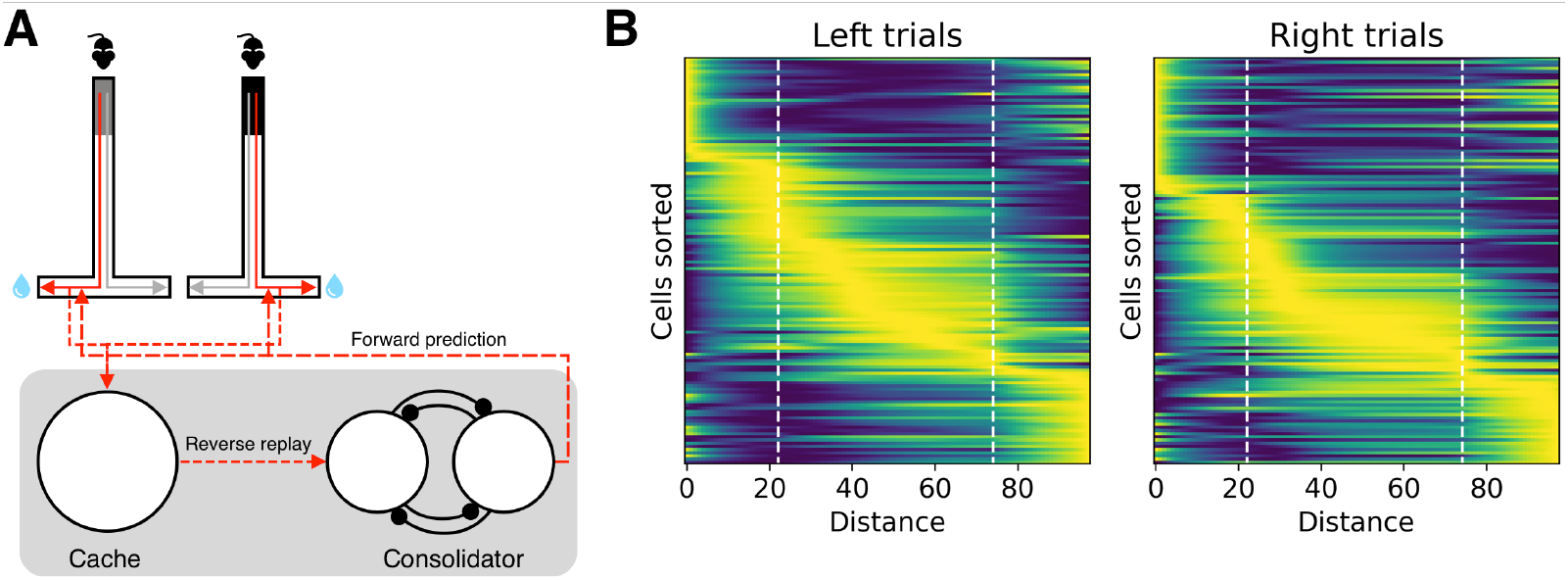
T-maze task trained with consolidator and cache. **A** Task structure and training paradigm. Upper: Animal makes a decision to run and then turn to left or right according to the cue type (black or grey block in the top of the **T**-maze) to get reward. Lower: The system first selectively receives rewarded sensory sequences (red trajectories in the T-maze) and stores it in the cache, which then performs reverse replay, providing a reversed input sequence to the consolidator. The consolidator then is trained to generate rewarding predictions. **B** Place representations in the hidden unit firing rates of the consolidator. For left and right rewarded trials, neurons are sorted according to the distance between the start point and positions with highest firing rates. The first vertical dashed line represents the distance at which the cue ends, while the second one indicates where the left/right decision point lies.

This was not intended to be a simulation of the brain itself. Rather, it was to test and examine the functionality of the model in a setting parallel to one commonly used to study memory in rodent models.

Briefly, the model alternately explored a T-maze and performed off-line learning after collecting enough experiences. As with rodents learning to complete the task, the model generated actions that directly impacted the sensory inputs that formed the episodic sequences. Thus, the learning task was two-fold - encoding the experienced sequences to enable accurate pre-diction of upcoming transitions and adaptive selection of actions to collect rewards. This is fundamentally different than the benchmark tests presented above in that the question is not whether the network can simply recall a training sequence.

Similar to the paradigm discussed previously, we performed another simulation to test both performance and a match with empirical data from the hippocampus. We set the sensory inputs to both consolidator and cache as a concatenated binary vector (*o*^*t*^, *a*^*t*^, *r*^*t*^) data stream at time step *t*, with *o*^*t*^ for visual observation, *a*^*t*^ for action and *r*^*t*^ for the presence of the reward.

After training, the {consolidator, cache} system successfully mastered the task with a correct performance rate above 90% (see Figure 8D). The performance rise is driven by a series of gradually decreasing replay epochs of training (Figure 8E). Besides, since this system requires training for both forward and backward circuits in the consolidator, the replay rate for both directions is balanced by definition (Figure 8F). These results account for the empirical finding that as the animal gets familiar with the task, the replay events occur less often [35] (Figures 8A, 8B and 8AC). To investigate the representation of tasks in the system, we sorted neurons’ normalized firing rates according to the distance between their positions with highest firing rate and the start point. Similar to some previous work [36, 37], the results (Figure 7B) show that place cells-like structure emerged after training, and some neurons are biased to crucial positions such as the end of the cue and the turning point.

**Figure 8:**
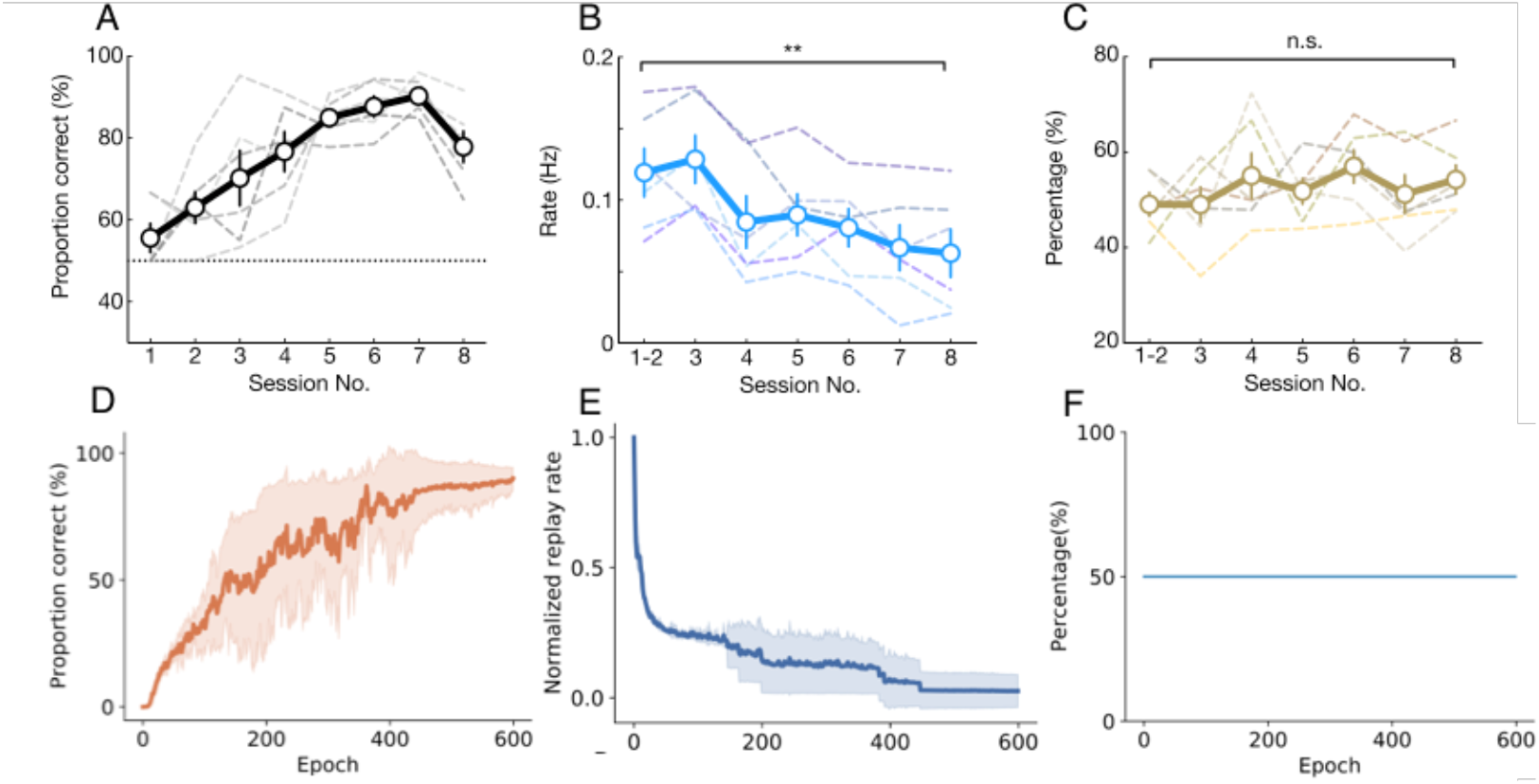
Model behavior in T-maze navigation task in comparison with results from [35]. Top row: **A, B** and **C** are adapted from Fig.4 in [35]. Bottom row: **D, E** and **F** are results from the the consolidator-cache model. **A**,**D:** Throughout training the model and animal performance both steadily increase. **B**,**E:** During the training process, the replay rate gradually decreases as the performance increases. **C**,**F: C** Animals have relatively balanced forward and reverse replay in the all training sessions regardless of performance. **F** In the system of the cache and consolidator, the replay rate is balanced by definition as forward replay involves learning of backward circuits and backward replay trains forward circuits.

## 4. Conclusion

In this article, a novel learning system, R2N2, is developed to address the long-standing question of biologically plausible learning of RNNSs. It is composed of two components: a fast RNN that stores and replays experiences (cache) and a statistical learning RNN (consolidator).

We have shown that R2N2 is capable of running itself in a reversed order and shows improved performance relative to other models based on recurrent weight updates computed in the backward phase in various tasks. In addition, by applying this model to a rat navigation task we showed its power as a whole in sequence learning and that it captures several observed phenomena in previous experiments such as balanced replay and place cell encodings.

## 5. Discussion

### 5.1. The R2N2 model

The massive success of deep learning models and their similarities with biological neural networks at both the behavior and neural dynamics level have captured the attention of the neuroscience community on many different questions, with one of the most crucial being how the full error gradient learning in RNNs might be implemented in the brain due to its recurrent (lateral connections) and hierarchical (multi-layer) connectivity nature.

To form episodic memories, brains need to encode protracted temporal sequences of states with high efficiency. RNNs can accomplish this task when trained with the biologically implausible machine learning algorithm BPTT. To build a biologically plausible alternative of BPTT, we divided the problem into three pieces and applied solutions to each to form a novel integrated learning system: R2N2. The first two pieces eliminate the need for an external record of the spatiotemporal activity pattern to perform offline learning. One piece establishes a way to reconstruct the input sequence activity in the backwards direction. This was solved with an RNN using a reciprocating weight structure. The second piece establishes a way to reconstruct the recurrent layer activity. This was solved in the consolidator network with a pair of competing, complementary subnetworks. Finally, the third piece eliminates the need for either weight transport or reversible synapses to back-propagate error. This is solved using feedback alignment.

Consequently, R2N2 consists of two major RNNs: 1) The consolidator, a main network that is to be trained and that serves as a long-term memory store for inference; and 2) The cache, an auxiliary network that supports the training of the consolidator by performing one-shot learning with the input layer activity sequences and thus providing training samples to the consolidator. Besides the constraint of only local information at synapses, as a seemingly obvious biological constraint in the brain, most of the work trying to solve this problem has focused on the online side [22, 13] as a workaround. Unlike its online alternatives to BPTT, R2N2’s underlying principle is that it has a backward phase to compute and assign credit to synapses in recurrent projections without violating the locality constraint.

The ability to compute the error feedback signal through numerous time steps may account for the advantage over previous localist supervised sequence learning models such as echo state network[33, 38] and RFLO [13], as it extends the error gradient farther back in time.

Since the consolidator is gradually trained to perform reverse replay in a nearly perfect way, we further speculate that the consolidator could, in turn, train other similar consolidator instances in the cortex in a ”bootstrapped” manner to implement distributed knowledge representation across distinct brain regions.

### 5.2. Biological Implications

Our model bears some similarity to the complementary learning systems (CLS) framework regarding the relative roles of the hippocampus and cortex. Typically the hippocampus is cast as the fast learner and the cortex the slower learner [39], and more recently the role of replays has been incorporated into the framework [40]. The R2N2 model suggests that the cache and consolidator functions (analogous to fast and statistical learning, respectively) may both be carried out within the hippocampal region, as well as between the hippocampus and neocortex. For example, the cache could be implemented in CA3 pyramidal neurons with recurrent lateral excitatory projections, which have the same arbitrary spatial association and pattern completion capability. With a trainer providing reversed sequence samples, the consolidator, which could be a circuit in the entorhinal cortex receiving inputs from CA3, could learn statistics in the data stream and solidify the short-term memory in the hippocampus to provide longer-lasting memories. When assembled together, this system could be triggered and tuned by reward-related signals as we did in the T-maze simulation to ensure the sequence being replayed and learned is rewarded and beneficial for the animal, which has been found to be the case in the hippocampus [27].

Recent work has similarly argued that both fast and statistical learning may take place within the hippocampus, with the entorhinal cortex to CA1 pathway providing statistical learning, and the pathway from CA1 to dentate gyrus to CA3 providing fast learning [41]. The R2N2 model is consistent with this anatomical delineation but does not exclude other possible functional mappings. Another implication provided by R2N2 is its utilization of the reverse replay phenomenon found in hippocampus [15, 3], which is the key element that drives the whole model to learn sequences. However, many of the existing models [42, 43] for reverse and forward replay do not account for sequence learning at all or have a limited learning capacity. They are limited in that those models are built on handcrafted attractor connectivity patterns and thus usually have only one or few spatially clustered neurons active at each moment, which is functionally equivalent to one-hot encodings and puts a limit (*N* =the number of neurons) on their learning capacity. Instead, the consolidator in our model builds connectivity matrices for the reverse replay of arbitrary neuronal activation pattern sequences without any prior assumptions on the spatial distribution of synapse strengths, which is far more flexible and biologically realistic considering the high dimensional nature of spiking activities in the brain. This also matches previous observations that the reverse replay in the brain is key for sequence learning [3, 4, 5, 6, 7, 8, 9]. It implies that the hippocampal cortical system may be a neural instantiation of BPTT, and our proposed model might account for the underlying mechanism of reverse replay as well as its computational role in learning.

A possible neural realization of this consolidator-cache system might be the entorhinal-hippocampus communication system. First, there is empirical evidence showing it is the backward running phase (reverse replay), rather than the forward running phase (forward replay), in the hippocampus during immobilization that is crucial for the animal’s later performance after experiencing the task environment [27]. Some other recent studies even show that prolonged reverse replay enhances task performance [8], while destroyed reverse replay leads to failures in task performance [9]. Second, R2N2 also suggests the importance of internal clock signals, as the consolidator and cache each oscillate to generate state updates. The importance of oscillating clock signals is consistent with several hippocampal cell types which show either the greatest or smallest activity levels at the peak of the theta cycle or a ripple [44]. Our simulation results also reveal that during learning the system shows similar characteristics and internal representations to place cells and replay rate effects observed in previous studies [36, 37, 35].

Together, these observations imply the existence of offline backward learning in recurrent neuronal networks, which is conceptually isomorphic with the temporal unfolding process in BPTT. However, there also exist some limitations in the current implementation of R2N2. For instance, the speed of forward replay and reverse replay is the same in our simulation while the reverse replay in the hippocampus is usually highly compressed in time compared with the forward running process [15].

The temporal symmetry of reverse replay in R2N2 is due to the same time constant we used in Eq.1 for both forward and reverse modes. In future research, we will explore addressing this issue by diving into the level of spiking networks, since the speed change is caused by the spike interval reduction, which can be implemented with a discrete version of the consolidator that preserves the symmetry but in which the timing can be freely tuned. The speed of network evolution may also be controlled by a clocking mechanism similar to a CPU clock, in which the frequency of an oscillatory signal as in Figure 6 may control the speed of the network.

### 5.3. Summary

In summary, this article provides a new possible approach for biological RNNs to learn sequential tasks: R2N2. R2N2 can memorize sequences in a one-shot way and transfer the experience to long-lasting synaptic changes through reverse replay. When it comes to cognitive tasks, the consolidator-cache system treats different types of tasks under a unified sequence prediction framework and solves it with rewards as a signal for reverse replay. The whole process is based on competitions between different synaptic projections, i.e., the competition between *f*_∗_ and *g*_∗_ in the consolidator and *W*_*O*_ and *W*_*E*_ in the cache, which does not require any non-local information or weight symmetries. When compared with other online alternatives to BPTT, R2N2 can better propagate the temporal information in the training phase and thus has better performances in some tasks. This computational superiority is driven by the use of reverse replay as an error propagating mechanism, which also raises several experimentally testable predictions for future research on sequence learning in the brain. First, an imbalanced synaptic projection (e.g., decreased excitatory level in one projection) between different neural assemblies may lead to impaired reverse replay since in our model the reverse replay in the consolidator relies on competition of different projections between neuronal groups. Second, as in the cache, an internally generated pseudo-periodic signal is responsible for the transition between firing pattern attractors, one may expect to see induced reverse replay with external periodic signals acting on the gating neurons for the CA3 network or corrupted reverse replay with aperiodic perturbations.

## 7. Acknowledgements

The authors thank Ehren Newman for extensive and helpful discussions and comments on the manuscript.

## 8. Data Availability

The code for the simulations in this paper can be found at https://github.com/CogControlLab/R2N2

## Appendix A. Hidden layer error computation

In our implementation, the specific process to update the postsynaptic component, 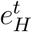, in Eq.3 is discretized as a difference equation from time *t* = 0 to *t* = *T*. Consider a simple case in which from (0, *T*) the task of the network is only to generate a proper output *O*_*T*_ at the last time step *t* = *T* and the overall loss function is then defined as 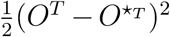 with 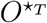 being the desired output. If neuronal group *B* is connected to the output layer *O* through a linear mapping *W*_*O*_, then we have

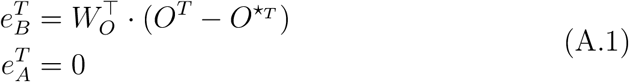

for the last time step.

For all other previous steps *t* ∈ (0, *T*−1), we have two coupling equations to compute the error signals for both *A* and *B* iteratively:

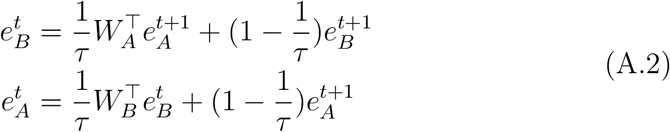

We include a leaky term in the the above update equations to maintain consistency with the leaky nature of the activity updates in Eq. 1.

With the help of Feedback alignment algorithm, the 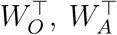 and 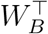 in the above equation can further be replaced by random fixed matrices *β*_*O*_, *β*_*A*_ and *β*_*B*_, for *t* = *T*, then we have:

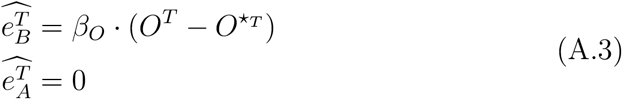

and for *t* ∈ (0, *T* − 1):

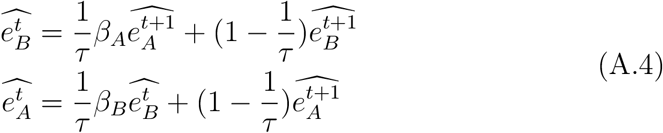

If we generalize the case of only generating output *O*^*T*^ at *T* to outputs at each time step *O*^1^, *O*^2^,, *O*^*T*^, the corresponding *e*_*A*_ and *e*_*B*_ will be similar with each time step to the neuronal group connected to the output layer receiving an extra non-zero output error term used in Eq. A.3

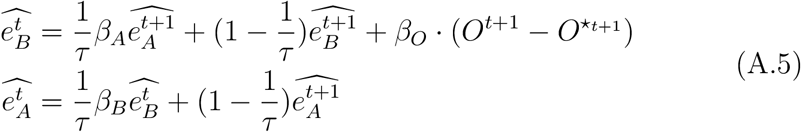

When connecting with the cache network, since we’re taking a sequence prediction paradigm, 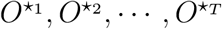 will be the activation sequence of the cache, which can perform reverse replay itself to provide the reversed desired output sequence required in Eq. A.5.

